# NMDA receptor blockade causes selective prefrontal disinhibition in a roving auditory oddball paradigm

**DOI:** 10.1101/133371

**Authors:** RE Rosch, R Auksztulewicz, PD Leung, KJ Friston, T Baldeweg

**Affiliations:** Wellcome Trust Centre for Neuroimaging, University College London, 12 Queen Square, London WC1N 3BG, UK; Developmental Neurosciences Programme, UCL Great Ormond Street Institute of Child Health, University College London, 30 Guilford Street, London WC1N 3EH, UK; Department of Psychiatry, University of Oxford, Warneford Hospital, Warneford Lane, Oxford OX3 7JX, UK

## Abstract

N-methyl-D-aspartate receptors (NMDARs) are expressed widely throughout the human cortex. Yet disturbances in NMDAR transmission – as implicated in patients with schizophrenia or pharmacologically induced – can cause a regionally specific set of electrophysiological effects. Here, we present a double-blind placebo-controlled study of the effects of the NMDAR blocker ketamine in human volunteers. We employ a marker of auditory learning and putative synaptic plasticity – the mismatch negativity – in a roving auditory oddball paradigm. Using recent advances in Bayesian modelling of group effects in dynamic causal modelling, we fit biophysically plausible network models of the auditory processing hierarchy to whole-scalp evoked response potential recordings. This allowed us to identify the regionally specific effects of ketamine in a distributed network of interacting cortical sources. Under placebo, our analysis replicated previous findings regarding the effects of stimulus repetition and deviance on connectivity within the auditory hierarchy. Crucially, we show that the effect of ketamine is best explained as a selective change in intrinsic inhibition, with a pronounced ketamine-induced reduction of inhibitory interneuron connectivity in frontal sources. These results are consistent with findings from invasive recordings in animal models exposed to NMDAR blockers, and provide evidence that inhibitory-interneuron specific NMDAR dysfunction may be sufficient to explain electrophysiological abnormalities of sensory learning induced by ketamine in human subjects.

Significance Statement
Dysfunction of N-methyl-D-aspartate receptors (NMDARs) has been implicated in a range of psychopathologies, yet mechanisms translating receptor-level abnormalities to whole-brain pathology remain unclear. We use computational modelling to infer microcircuit mechanisms by which ketamine, an NMDAR-blocker, alters brain responses to changing sequences of sounds that rely on sensory learning. This dynamic causal modelling (DCM) approach shows that ketamine-effects can be explained with brain region specific changes in inhibitory interneuron coupling alone, with a striking reduction in inhibition in the prefrontal cortex. This suggests that NMDAR-effects on excitation-inhibition balance differ between brain regions, and provides evidence from healthy human subjects that NMDAR blockade may cause prefrontal cortex disinhibition, one of the mechanisms hypothesised to underlie psychopathology in schizophrenia.

## INTRODUCTION

N-methyl-D-aspartate receptor (NMDAR) hypofunction is believed to be one of the primary causes of schizophrenia, a common neuropsychiatric condition (1–5). In healthy human subjects, pharmacological interventions such as exposure to ketamine, which acts as a non-competitive NMDAR antagonist, can in part reproduce a recognisable set of symptoms and signs (such as delusional beliefs, hallucinations, working memory deficits and social withdrawal), and electrophysiological brain abnormalities associated with schizophrenia (6–8). Molecular genetics and cellular neuroscience have provided many important insightsinto the underlying pathophysiology. These approaches often identify abnormalities at the level of neurotransmitters and their receptors (9, 10), neuronal microstructural changes (11, 12) and localised features of neuroanatomy (13). However, relating pathophysiological hypotheses based on these findings to dysfunction at the whole-brain level in schizophrenia remains challenging (1, 14).

Computational models of neuronal function – often framed at the scale of neuronal populations – offer a bridge between putative synaptic mechanisms of disease and observed psychopathology orendophenotypes: they allow both *bottom up* predictions of the effects of synaptic changes on the function of neuronal ensembles and networks (15, 16), and (through model inversion techniques) a *top down* inference about plausible pathophysiological mechanisms underlying observed whole-brain phenotypes (17–19).

The auditory mismatch negativity (MMN) is generally impaired in schizophrenia (17, 20, 21) and a similar impairment can be induced with ketamine in healthy human subjects. The MMN is based on a stereotyped response to deviants or oddballs embedded within regular stimulus sequences, and has emerged as an important tool for probing brain function using non-invasive electroencephalography (EEG) recordings in human subjects. MMNs are difference waves of evoked response potentials (ERPs) to the unexpected *deviant* stimuli compared to repeated *standard* stimuli (22–28). This difference can be observed in a variety of paradigms, ranging from deviant responses in simple auditory, visual or somatosensory stimulus sequences to MMN-like responses to violations of syntactic or semantic expectations in language paradigms (29–31).

The domain-general nature of MMN responses suggest that they arise from general mechanisms underlying sensory processing in the brain. A dominant theory that accommodates such perceptual inference is formalised in the predictive coding framework (32, 33): Based on Helmholtz’s notion that the brain attempts to infer the causes of sensations (34, 35), predictive coding proposes that the brain generates predictions of its sensory input. When sensation deviates from these predictions, prediction error signals are generated and passed along the sensory processing hierarchy. This framework integrates competing views regarding the neurobiological basis of MMN responses. According to the predictive coding account, the MMN arises both from disruptions of neural adaptation (classically regarded the *neural adaptation hypothesis*, (36)), and adjustments of the model on which predictions of future stimuli are based (classically regarded the *model-adjustment hypothesis*, (37)). There is increasing evidence across different experiments, sensory domains and even species suggesting that predictive coding provides a good explanation of MMN type responses (28, 33, 38–40). However, currently there is limited evidence on how the physiological mechanisms underlying the MMN are affected by ketamine.

One empirical approach to test hypotheses regarding the neuronal processes underlying the MMN is dynamic causal modelling (DCM). In DCM of ERPs, biophysically informed models of cortical microcircuitry (41, 42) – typically based on neural masses – are fitted to measured data using variational Bayesian model inversion techniques (43, 44). This approach has been widely applied to model changes in neuronal coupling that underlie observed ERPs in auditory MMN paradigms (17, 22, 27, 45–48), producing network-wide electrophysiological consequences, during sequence learning (49) and responses to deviants (46, 50) that are consistent with predictive coding. The DCM approach has previously been applied both to MMN responses in patients with schizophrenia (48), patients with psychosis and family members (17), and healthy human subjects treated with ketamine (51), all of which have identified modulations of connectivity between cortical sources as underlying the observed MMN differences between groups. The latter study specifically identified a single forward connection from left primary auditory cortex to left superior temporal gyrus to be altered significantly by ketamine, by applying classical statistics to the connectivity estimates provided by DCM.

Here, we build on the existing literature by applying recent hierarchical Bayesian modelling procedures to a double-blind placebo-controlled auditory MMN study of the effects of ketamine on coupling in the auditory hierarchy. Using a parametric empirical Bayes approach to group inversion of subject-specific DCMs (52), we use a hierarchical model that includes (i) within-session coupling changes explaining ERPs to both deviant stimuli (i.e. modelling the classical mismatch response) and to repetitions of the same standard (i.e. modelling repetition suppression effects); and (ii) within-subject, but between-session differences in the first-level DCMs induced by ketamine in the double-blind cross-over trial design.

In the original description of DCM analysis of MMN responses (22, 45, 46), parameter changes are typically restricted to synaptic coupling changes that are plausibly affected by sensory input and condition specific changes. Therefore previous DCM analyses of ketamine effects (51) limit the possible explanations of observed ketamine effects to a subset of cortical coupling. However, pharmacological interventions can potentially have a more distributed effect on cortical function; by influencing the excitation/inhibition balance within cortical microcircuits, by changing the intrinsic timescales of cortical areas, or by modulating postsynaptic gain (53). These effects can now be modelled efficiently with a hierarchical model that distinguishes between within session (e.g. oddball) and between session (e.g., ketamine) effects. In this work we used recent developments in hierarchical modelling (i.e., empirical Bayes) and Bayesian model reduction to identify the best (reduced) model that explains both oddball and ketamine effects. Bayesian model reduction identifies the best model of treatment effects that could potentially be expressed in an unknown combination of parameters or connections (52, 54). It does this by removing redundant parameters; thereby increasing model evidence. We used Bayesian model reduction to identify which combination of intrinsic (within-source) or extrinsic (between-source) synaptic connections changed during the presentation of oddball stimuli and how these changes were contextualised by the administration of ketamine.

We hoped to replicate previous findings regarding network-wide changes underlying MMN responses in the brain, and to identify the network coupling changes underlying short term sensory learning during repetitions of the same stimuli. Furthermore, we want to characterise the effects of ketamine – as an NMDAR blocking agent – on the MMN response in healthy subjects. NMDA receptors are prevalent in the supragranular layers of the cortex, suggesting particular relevance of NMDAR transmission to backward connections in the cortical hierarchy that target superficial layers (55). However, NMDAR are unevenly distributed across cortical interneuron subtypes, indicating that the overall effects of NMDAR-blockade may be better represented in regionally-specific intrinsic coupling changes (excitatory, or inhibitory), that affect specific sub-populations rather than extrinsic coupling between regions (56).

ERP abnormalities – induced by NMDAR-blockade with ketamine – may be broadly underwritten by two distinct mechanisms: (i) Ketamine may alter the basic setup of the auditory processing network in terms of synaptic function (i.e. excitatory/inhibitory coupling, synaptic time constants, between-source connectivity) irrespective of the auditory context. Note that because of the nonlinearities in the neuronal system, even a context-invariant change in connectivity (i.e., in parameter space) may produce differential effects in standard versus deviant responses in the various ERPs (i.e., in measurement space). (ii) Alternatively, ketamine and the ensuing NMDAR-blockade may have a direct impact on synaptic plasticity induced by the sensory learning during the roving MMN paradigm. In other words, changes in measured deviant responses may be caused by ketamine sensitive plasticity; namely, an interaction between deviant and drug effects at the level of connectivity. Given that DCM estimates both synaptic plasticity induced by deviants and changes induced by ketamine, we can disambiguate these hypotheses based on measurements of MMN responses.

## RESULTS

### ERP results in sensor space

In this roving oddball paradigm, sequences of the same sound were repeated up to 36 times, before a new sequence began at another random tone frequency. This means that when averaging ERPs to the same position within sequences, each mean ERP contains responses to a range of physical tone frequencies – thus the averages differ not in physical property of the sound played, but only in their location within a sequence. Here, we show the resulting grand mean ERPs across all eighteen participants (details in Table 1) at the fronto-central (*Fz*) electrode for the 1st (deviant, D1), 2nd, 6th and 36th (standards, S2, S6, S36) exposures to the same sound as grand mean averages across subjects; separately for placebo and ketamine conditions (Fig 1A). Response to the first stimulus (D1) constitutes a typical deviance response, with an early negativity (N1, peak around 150ms), and later positivity (P3a, peak around 250ms) that differs significantly from the S36 ERP around those time intervals (Fig 1A: Deviance effect (red) paired t-test at each time point, *p* < 0.05, Bonferroni-corrected for multiple comparisons). Responses to the standard tones (S2, S6, S36) illustrate the build-up of a positivity peaking at around 120ms, known as memory trace formation. This positivity is significantly different between S2 and S36 for segments ranging from approx. 50ms to approx. 200ms (Fig. 2A: Repetition effect (green) – paired t-test at each time point, *p* < 0.05, Bonferroni-corrected for multiple comparisons). Ketamine reduced both deviance and repetition effects, compared to placebo with a collapse of the time windows for which these effects were significant (duration of significant deviance effect: placebo 112ms, ketamine 90ms; duration of significant repetition effect: placebo 120ms, ketamine 98ms). The difference, mismatch negativity waveforms (Fig 1B) peaks at a significantly higher amplitude for the placebo condition for the early (N1, paired t-test, (17) = 1.85: *p* = 0.04) but not the late (P3a, (17) = 1.10: *p* > 0.05) component. The ketamine-induced attenuation of the MMN is apparent across the whole scalp when plotting all channels (Fig 1C).

**Figure 1.**
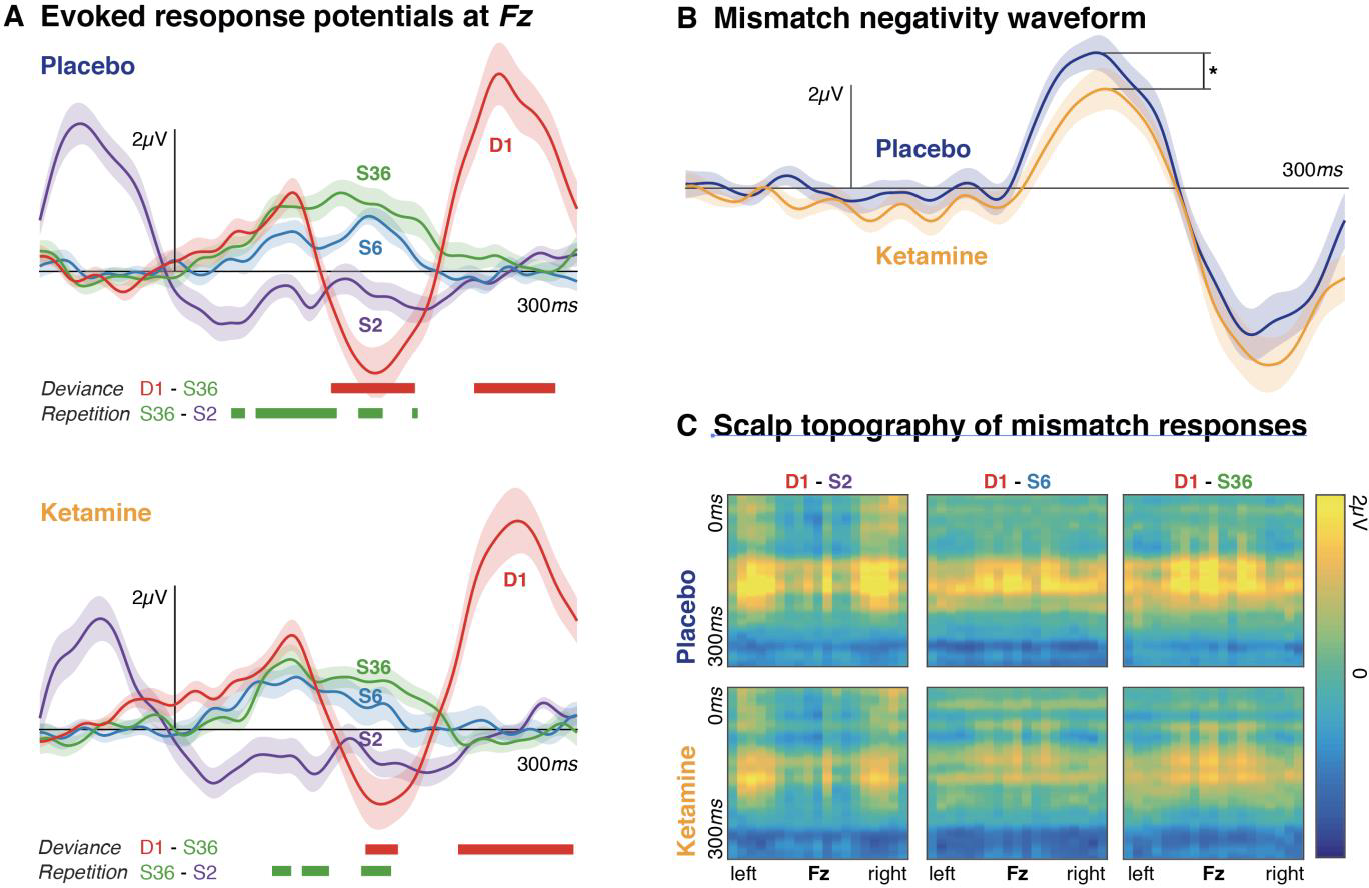
Ketamine causes a reduction in the mismatch negativity. (A) Evoked response potentials (ERPs) are shown for repetitions of a sound within a roving oddball paradigm. The first exposure to a sound within a sequence, *D*1, provokes a typical deviance response at the Fz electrode. ERPs for three different repetitions, *S*2, *S*6 and *S*36 show increasing positivity with a peak at approximately 120ms. The bold red lines indicate time points for which the S36 and D1 ERPs are significantly different across the group (i.e. the effect of deviance,‥ <0.05, Bonferroni-corrected for multiple comparisons); bold green lines indicate time points for which *S*36 and *S*2 are significantly different (i.e. the effect of repetition,‥ <0.05, Bonferroni-corrected for multiple comparisons, differences only tested for the 0-300ms peristimulus time interval). Ketamine reduces both the deviance and repetition effects. (B) Difference waveforms at Fz are shown for *D*1-*S*36. The peak amplitude of the around 150ms is significantly bigger for the placebo condition compared to ketamine. (C) The panels show the difference between *D*1 and *S*2, *S*6 and *S*36 respectively across time (y-axis), and channels (x-axis, arranged from left to right). Across all condition, there is a ketamine related reduction in mismatch responses.

**Table 1:**
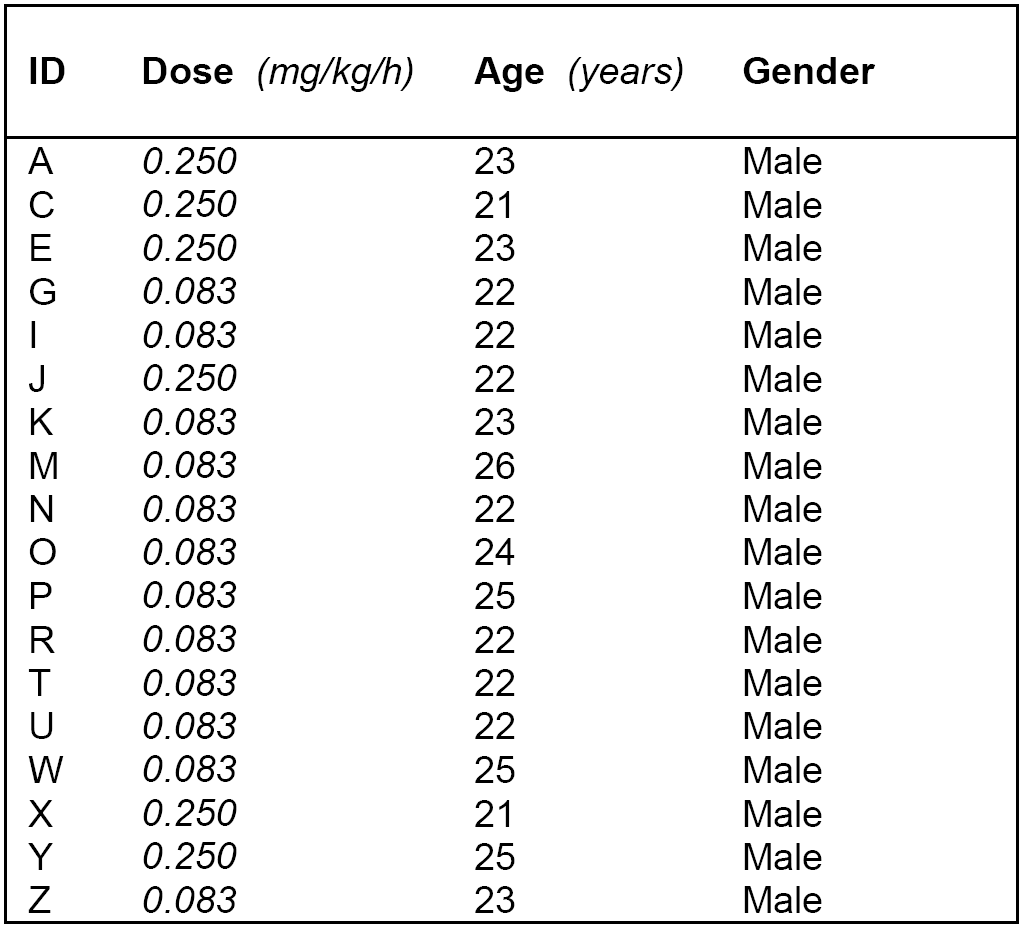
Study subject details.

| ID | Dose (mg/kg/h) | Age (years) | Gender |
| --- | --- | --- | --- |
| A | 0.250 | 23 | Male |
| C | 0.250 | 21 | Male |
| E | 0.250 | 23 | Male |
| G | 0.083 | 22 | Male |
| I | 0.083 | 22 | Male |
| J | 0.250 | 22 | Male |
| K | 0.083 | 23 | Male |
| M | 0.083 | 26 | Male |
| N | 0.083 | 22 | Male |
| O | 0.083 | 24 | Male |
| P | 0.083 | 25 | Male |
| R | 0.083 | 22 | Male |
| T | 0.083 | 22 | Male |
| U | 0.083 | 22 | Male |
| W | 0.083 | 25 | Male |
| X | 0.250 | 21 | Male |
| Y | 0.250 | 25 | Male |
| Z | 0.083 | 23 | Male |

### Effects of repetition on connectivity

Reptition effects were modelled as changes in connectivity parameters in a distributed cortical auditory network comprising three bilateral sources arranged along a processing hierarchy. Repetition effects or plasticity were modelled as a linear mixture of two time courses (shown in Fig 2A): a monophasic decay and a phasic repetition effect. This constitutes the full model; i.e., where both monophasic, and phasic repetition effects could modulate forward and backward connections as well as intrinsic modulations at each level of the cortical hierarchy (model *FBi* in Fig 2A). The model fits for this model in each subject are shown in Fig 3A).

**Figure 2.**
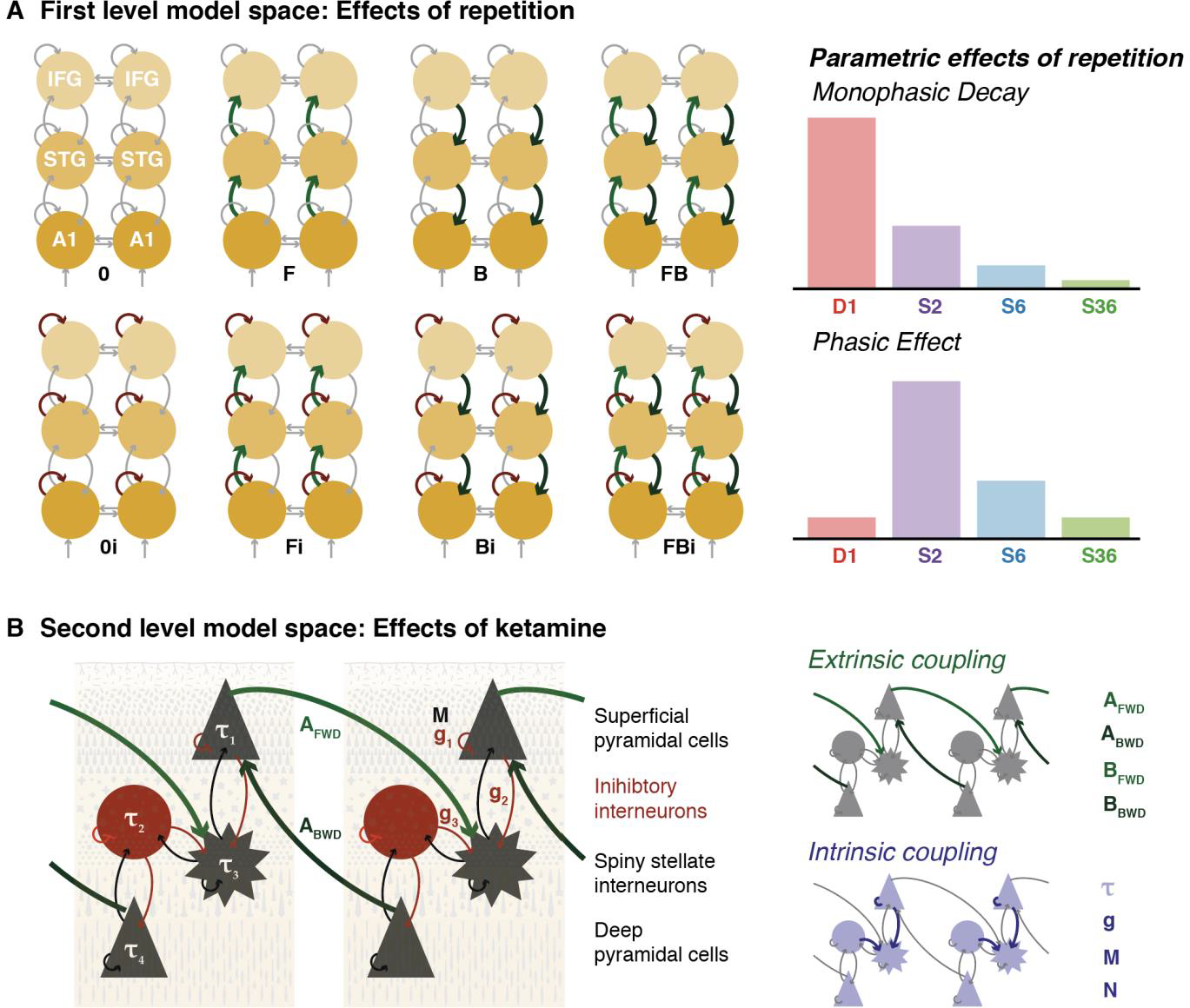
Two-level hierarchical DCM model space. (A) Effects of repetition were modelled as changes in a combination of forward/backward connections and intrinsic modulatory gain parameters in a coupled network of six cortical sources. Repetition effects were parameterised as a mixture of monophasic decay and phasic activation as shown in the right panels. (B) Ketamine effects were modelled at the second level; i.e. as systematic differences in connectivity parameters. Each cortical source corresponds to a canonical microcircuit, with four populations that is parameterised with extrinsic and intrinsic coupling parameters. The second level model space consists of combinations of these parameters which explain the differences between placebo and ketamine conditions. *A1 – primary auditory cortex; STG – superior temporal gyrus, IFG – inferior frontal gyrus*

Subsequently a set of reduced models was estimated using Bayesian model reduction. These reduced models comprised each of the models in Fig 2A paired with the monophasic decay effect, the phasic repetition effect, or both, resulting in a total of 24 models. Bayesian model comparison provides decisive evidence for the full model (i.e. *FBi* with both monophasic and phasic effects) at the group level (Fig 2B) and for each individual subject.

Bayesian parameter averages for forward connections, backward connections, and modulatory self-connections (shown here for A1 where the strongest effects were observed) are shown in Fig 3C, suggesting different time courses of changes for different types of connections. There is an overall reduction in forward and backward connectivity across repetitions. However, the largest change is earlier in the forward connections (D1 to S2) than backward connections (S2 to S6). Modulatory gain parameters are reduced from D1 to S2, before increasing between S2 and S36. These differences in the relative modulation of different types of connections underlie the differences observed in the model predictions for ERPs for different repetitions.

**Figure 3.**
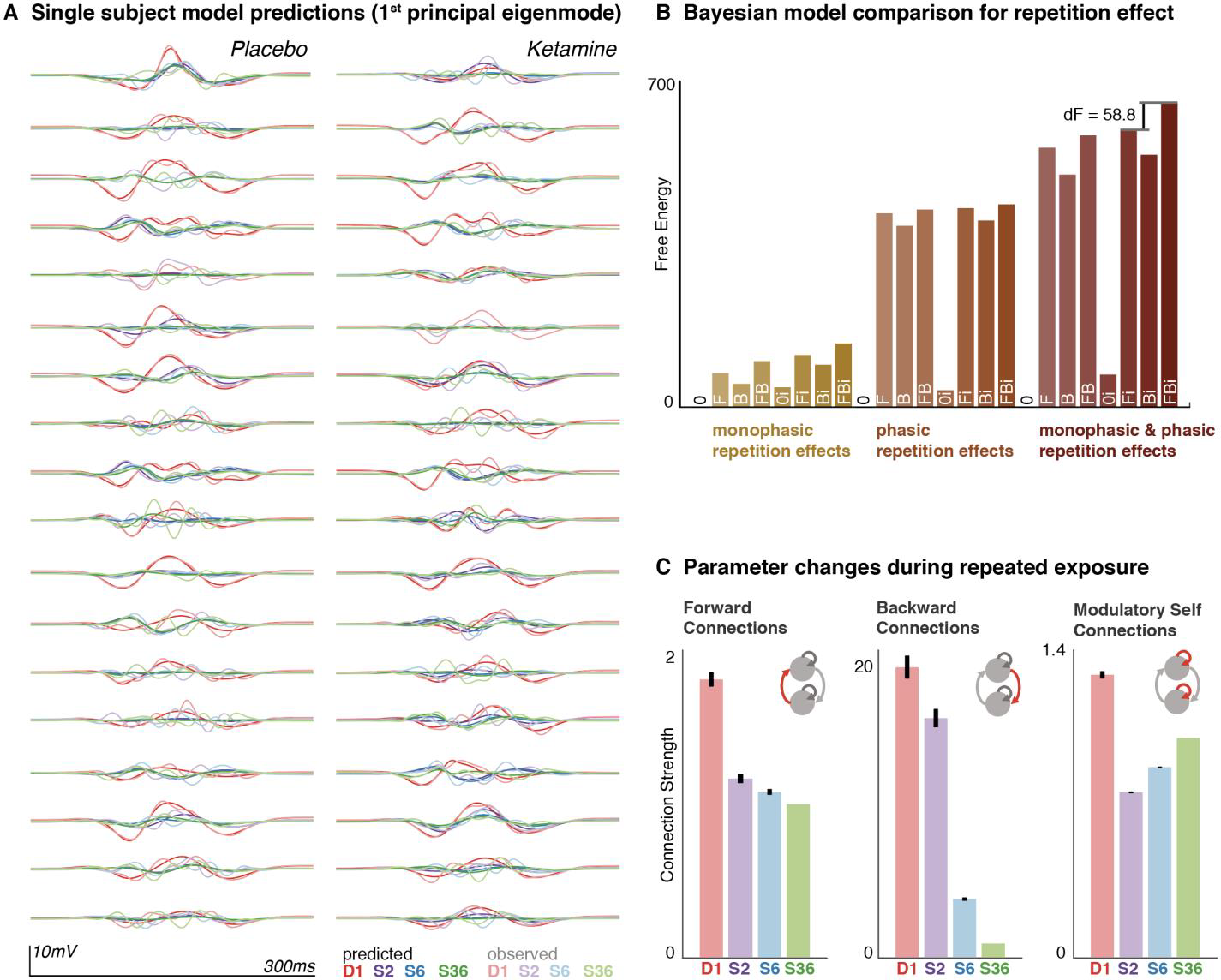
Repetition effects. (A) ERPs to the 1st (Deviant, D1), 2nd, 6th, and 36th (Standards, S2, S6, S36) presentation of a sound within a sequence were modelled in subject-specific DCMs. The first principal eigenmode of the prediction in sensor space (bold colours), and the corresponding mode of the empirical scalp data (light colours) are shown for each individual. These suggest a good fit for the main components of the ERP waves. **(B)** Bayesian model comparison was performed to compare models in which the repetition effect was monophasic, phasic or both, and included modulations of forward *F*, backward *B*, or intrinsic *i* connections and their combinations. The winning model across the group was the full model, where monophasic and phasic repetition effects impact on forward, backward and intrinsic connection. **(C)** Bayesian parameter averages acrossthis full model for each individual subject show changes in connection strength across repetitions for forward, backward and intrinsic modulatory connections. Error bars indicate the 95% Bayesian confidence interval.

### Effects of ketamine on model parameters

At the second level of the analysis, we combined all individual subject DCMs for placebo and ketamine conditions into a single PEB model to identify parameter changes induced by ketamine. Initially, we performed Bayesian model comparison across alternative second-level models where subsets of extrinsic or intrinsic model connectivity parameters (as detailed in Table 2) were free to explain the ketamine effect (Fig 4A). Across this model space, there is very strong evidence for modulations in, and only in the intrinsic connectivity (*g* parameters) within cortical sources. In other words, these and only these parameters are required to explain the ketamine induced ERP changes described above.

**Table 2:** Canonical microcircuit model parameters.

| Intrinsic coupling parameters |  | Extrinsic model parameters |  |
| --- | --- | --- | --- |
| $\tau_1$ | Superficial pyramidal cell (**sp**) time constant | $A_{fwd}$ | Extrinsic forward connection (**sp** to **ss**) |
| $\tau_2$ | Inhibitory interneuron (**ii**) time constant | $A_{bwd}$ | Extrinsic backward connection (**dp** to **sp**) |
| $\tau_3$ | Spiny stellate interneuron (**ss**) time constant | $B_{fwd}$ | Repetition effects on extrinsic forward connection |
| $\tau_4$ | Deep pyramidal cell (**dp**) time constant | $B_{bwd}$ | Repetition effects on extrinsic backward connection |
| $g_1$ | **sp** modulatory self inhibition | | |
| $g_2$ | **sp** to **ss** inhibitory connection | | |
| $g_3$ | **ii** to **ss** inhibitory connection | | |
| $M$ | Modulatory gain (**sp** self inhibition) | | |
| $N$ | Repetition effects on modulatory gain | | |

**Figure 4.**
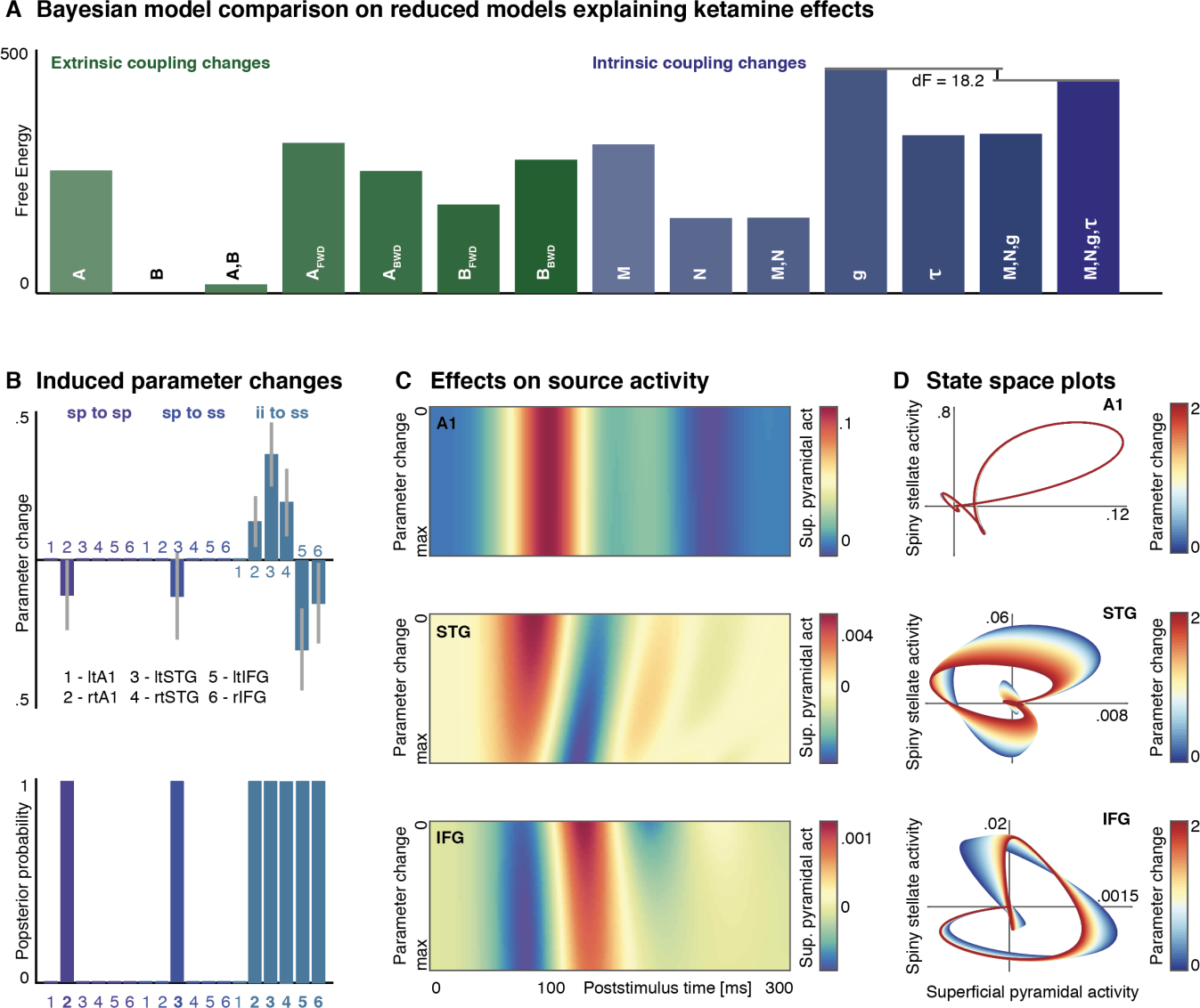
Ketamine causes frontal lobe disinhibition. **(A)** Using PEB, 14 alternative second level models were considered, explainingdifferences between ketamine and placebo with changes in combinations of parameters. Bayesian model reduction shows that the model with changes in intrinsic connection parameters (*G*) best explains the effects of ketamine on the ERPs. **(B)** Estimated parameter changes with Bayesian 95% confidence intervals (top), and posterior probability of the parameter being affected by ketamine (bottom) are shown. Significant changes were only observed in a subset of *G* parameters, with the largest effects estimated for inhibitory interneuron (*ii*) connections to spiny stellate (*ss*) cells. In the bilateral STG, there was an increase in *ii* inhibition on *ss*, while in the bilateral IFG there is a ketamine-induced disinhibition of *ss*. **(C)** The effects of opposing changes in *ii* to *ss* inhibition at different hierarchical levels are shown in source space. Each graph shows superficial pyramidal cell (*sp*) activity in different regions for the 0 – 300ms poststimulus interval with concurrent, but opposite modulation of parameter the *ii* to *ss* inhibition: In the STG the (log-scaled) connection strength is increased from 0 to 2, whilst in the IFG the strength is decreased from 0 to-2. This modulation causes an attenuation and increase in latency in the IFG response, with concurrent attenuation of early STG responses and a decrease in the latency of the response. **(D)** Neuronal state space plots show the relationship between *sp* and *ss* activity for different hierarchical levels and for increasing changes to the *ii* to *ss* inhibition. There is minimal effect on the A1. For STG, the parameter changes induce a reduction in *ss* response amplitude compared to *sp* and an overall shift towards more negativepopulation output. In the IFG there is an inverse reduction of *sp* response amplitude compared to *ss*.

An inspection of the winning second-level model revealed that of these *g* parameters, only a subset is affected by ketamine (Fig 4B). The biggest effect size is seen in *g*_*3*_ that represents the strength of inhibition supplied by inhibitory interneurons to excitatory spiny stellate cells. This parameter is modulated in opposite direction in the lower areas of the hierarchy (increased in right primary auditory cortex *A1*, left and right superior temporal gyrus *STG*; decreased in left and right inferior frontal gyrus *IFG*).

To simulate the (highly nonlinear) effects of these parameter changes on observed ERPs, we implemented a forward model based on the grand mean DCM inversion. Starting from the grand mean parameter estimates, this simulation gradually increased *g*_*3*_ in bilateral STG, while decreasing the same parameter in bilateral IFG. The effects of these reciprocal changes on source-space ERPs are shown in Fig 4C. This analysis reveals an attenuation and small increase in latency in the IFG response that resembles the observed changes in the mismatch negativity response at the *Fz* electrode in Fig 2B. The responses at STG level are overall reduced in amplitude with a decrease in response latency. Very little effect is observed on responses in A1.

Further analysis of the relationship between the excitatory interneurons (spiny stellate cells) and the main output neurons (superficial pyramidal cells) is shown in Fig 4D. Plotted terms of estimated neuronal responses (i.e. source space), these graphs represent the evolution of spiny stellate and superficial pyramidal cell responses during the deviance ERPs, starting from baseline (where spiny stellate and superficial pyramidal cell activity is equal to zero), and returning to baseline as a cycle through neuronal state space. These plots show the relative impact of the *g*_*3*_ parameter changes on spiny stellate and superficial pyramidal cell-populations. Strikingly, in IFG – where *g*_*3*_, or the inhibitory interneuron inhibition on spiny stellate cells, is reduced – this plot reveals a decrease in superficial pyramidal cell amplitude with relative preservation of spiny stellate cell activity. Conversely, in STG where *g*_*3*_ is increased, the amplitude of spiny stellate cells is relatively decreased compared to superficial pyramidal cells, with an overall shift into more negative superficial pyramidal cell responses.

## DISCUSSION

Here, we present a double-blind placebo-controlled study of ketamine effects on cortical responses to disruptions and repetitions within regular auditory sequences. Using a hierarchical dynamic causal model, we replicate previous findings in regards to network changes that explain the response both to deviants and to an increasing number of repetitions of the same sound. Furthermore we identify region-specific changes in cortical microcircuits that may underlie the ketamine-induced changes in MMN. The modelling analysis here thus integrates short-term sensory learning effects, the different cortical responses induced by deviants and changes to the system caused by NMDAR-blockade with ketamine.

## Computational modelling links whole brain observations with synaptic mechanisms

The modelling presented here applies recent developments in the Bayesian treatment of dynamic causal models to questions that bridge conceptual scales: How does a pharmacological agent, whose action is reasonably well understood at the microscale of the individual neuronal membrane, produce observable brain phenotypes? How does systemic application of ketamine produce effects on specific evoked response potentials with regionally distinct patterns?

The neural mass models applied in this analysis are mildly nonlinear approximations of neuronal populations that effectively capture a wide variety of normal (57, 58) and abnormal (59, 60) brain states and responses. The Bayesian approach allows one to assess how well a given model or hypothesis explains a set of observations. This *mesoscale* modelling thus allows the testing of mechanistic hypotheses about non-invasive (human) data and therefore allows inference on hidden neuronal states in human subjects that cannot be directly measured (Moran et al., 2011).

One of the key insights from this approach is not whether a given model is *true* or *false*, but rather a direct test of whether a specific mechanistic hypothesis is *sufficient* to explain a complex set of observations. Here, we compared a range of possible ketamine effects – on extrinsic excitatory coupling, intrinsic inhibition, and/or postsynaptic gain parameters – against each other based on whole-scalp ERP data from several experimental conditions at a subject-specific level. The modelling allowed us to identify those parameters that have most explanatory power, and compare them with existing literature from different experiments. For example, although we allowed for an effect of ketamine on plasticity (i.e. changes in connectivity as opposed to connectivity *per se*) this effect was a redundant component of our models – and was eliminated during Bayesian model reduction. In other words, a more parsimonious explanation for the effects of ketamine on oddball or mismatch responses is a change in the excitability (or disinhibition) of particular sources within the auditory hierarchy. This effect contextualises repetition dependent changes in a way that is sufficient to explain the effect of ketamine on the mismatch negativity.

Despite the complexity of both the datasets and the network models used for the model inversion, the results suggest that (i) acute NMDAR blockade effects on MMN responses can be explained by effects on a small set of key parameters, and (ii) the parameters identified are consistent with a wealth of findings from the basic science literature and studies of humans exposed to ketamine and patients living with schizophrenia. This non-trivial link between MMN changes and intrinsic inhibition within frontal microcircuits could not have been made without explicit computational modelling of human EEG generators, and provides further evidence for the relevance of GABAergic interneuron specific NMDAR hypofunction in the pathogenesis in psychotic illnesses such as schizophrenia (61, 62) – this instance, directly from a placebo-controlled human experiment.

## Deviance responses are caused by network-wide connectivity changes

There is ongoing debate regarding the computational and neuronal basis of mismatch negativity responses in the human brain. These are often broadly summarised in terms of (i) the *neural adaptation hypothesis*, according to which MMNs can be explained by bottom-up dishabituation responses of neuronal populations that adapt to auditory features of repeated stimuli (36); and (ii) the *model adjustment hypothesis*, according to which MMNs represent an error detection signal that result from a disruption of learned regularity in an auditory context (37).

The existing literature suggests that both changes in primary auditory cortex sensory gain (consistent with the *neural adaptation* perspective), and forward/backward changes in extrinsic regional coupling (consistent with the *model adjustment* perspective) are required to explain the difference between whole scalp responses to standards and deviants (22, 46, 47). The results of the placebo component of this study support the same conclusion: Both changes in the intrinsic modulatory gain and the extrinsic coupling between sources is required to best explain the differences between deviance responses and responses to standard sounds.

These findings are in keeping with a predictive coding account of MMN generation, aiming to integrate *neural adaptation* and *model adjustment* perspectives. According to the predictive coding framework, the brain actively generates predictions of the environment that are passed down from higher areas of the sensory processing hierarchies. These predictions are compared with incoming sensory input to produce prediction error signals, when sensation does not match the prediction. This prediction error signal passes up the cortical hierarchy, whilst at the same time resulting in a readjustment of intrinsic gain, or post-synaptic sensitivity in the primary sensory cortical areas (35, 40).

## Sensory learning causes distinct patterns of change for different coupling parameters

Using long sequences of sound repetitions allowed quantitative modelling of repetition responses: In the roving oddball paradigm, a deviant sound is repeated again and again until it becomes the new standard tone – and does not elicit a deviance response. Previous DCM analyses on repetition-induced changes in network coupling parameters have identified distinct temporal patterns in intrinsic *vs.* Extrinsic coupling changes: using short auditory sequences, Garrido et al (49) identified both monophasic decay and phasic change components in connectivity over repeated stimulus presentations. However, they found a clear difference in extrinsic connections, which were consistently reduced with each repetition; and intrinsic connections, which showed an initial phasic decrease before slowly increasing with repetition.

These findings are replicated in the independent dataset analysed here: extrinsic connectivity consistently decreases with repetition, whilst intrinsic connectivity parameters at the primary auditory cortex show a first dramatic decrease, before slowly increasing with repetition. Interestingly, our findings also reveal a temporal dissociation between forward and backward connections: while forward connection strengths return to their baseline value almost immediately (i.e. at the 2nd exposure to the sound), backward connection strengths remain higher for longer, with similar, persistently high parameter values at 1st and 2nd exposure to the sound.

This asymmetry in the time course of forward and backward plasticity revealed by the paradigm used here reflects a more general difference in the temporal dynamics along the cortical hierarchy. Primate cortical areas are hierarchically ordered; not only in terms of functional processes, but also in the timescales of cortical activity (63, 64), with time constants that can differ by orders of magnitude. These distinct temporal scales in a hierarchically structured network support efficient tracking and prediction of complex sensory input (65), through the hierarchically segregated tracking of fast and slow changes at different levels of the hierarchy. Distinct temporal scales may help integrating auditory objects within auditory contexts, purported as one of the functions of stimulus-specific adaptation, a marker of auditory learning within primary auditory cortex (66, 67).

Our findings speak to this hierarchical separation in time scales in the learning of sensory regularities. After the deviant, transiently increased forward (bottom-up) connections return very quickly to the baseline value, consistent with the short time-constants in lower cortical areas and the AMPA-predominance in these connections. Changes in backward (top-down) connections remain increased for a larger number of repetitions and as such encode synaptically the preceding sensory context (i.e. the recent occurrence of the deviant).

## NMDAR blockade has regionally specific effects on intrinsic connectivity

Using a hierarchical Bayesian model of both repetition and ketamine effects means that all available data (i.e. individual subjects’ whole-scalp ERPs to the four types of tones, D1, S2, S6 and S26) are used to inform model comparison and the estimation of the effect sizes. Furthermore, Bayesian model reduction allowed for data-driven reduction of the model complexity to identify the most relevant model parameters affected by the ketamine intervention. Notably, our findings broadly replicate those presented by Schmidt et al. (51) – if we restrict our second level model space to only those parameters considered in the standard DCM analysis (i.e. A, B, M, and N parameters, cf. Fig. 4A), we can replicate their findings. Among those models, the highest model evidence is provided by a model where ketamine affects the forward connections between cortical sources, an effect also described in their analysis. However, in a larger model space, the regional effects of ketamine can be better explained with a different set of parameters.

Our approach suggests that changes in a limited set of regional intrinsic connections best explain the ketamine effects. Furthermore, the most significant changes affect the same connection type: the inhibitory connection from inhibitory interneurons to spiny stellate interneurons, which is one of the coupling parameters between superficial (spiny stellate and superficial pyramidal cells) and deep neuronal oscillator pairs (deep pyramidal cells and inhibitory interneurons), supporting fast and slow intrinsic dynamics respectively (68). The direction of the modulation of this parameter depends on the affected cortical sources: the parameter estimates suggest that ketamine (assumed to be best described as NMDA receptor blockade) causes a decrease in inhibitory interneuron to spiny stellate inhibition in STG, with a concurrent increase in inhibitory interneuron to spiny stellate inhibition for IFG. When simulating the effects of these parameter changes at the different layers of the hierarchy, the IFG disinhibition notably results in a mild relative reduction in superficial pyramidal cell amplitude, whilst the STG response is characterised by larger amplitude reductions for both spiny stellate cell and superficial pyramidal cell populations.

These findings speak to the complexity of NMDAR-based transmission within cortical circuits. NMDA receptors have distinct distributions in different neuronal populations and exert direct effects on pyramidal cells, excitatory and inhibitory interneurons as well as modulating GABAergic transmission (56). We have focused on the auditory system and the mismatch negativity paradigm. Dynamic causal modelling of ketamine effects in other systems also implicates prefrontal regions; e.g. in a DCM study of fronto parietal coupling under ketamine (69) and in a study of fronto-hippocampal coupling (70). Interestingly, cell-type specific deletion of NMDAR on parvalbumin positive fast inhibitory interneurons are already used as a preclinical model of schizophrenia in a interneuron subtype specific mouse NMDAR knock out (71) and even result in some further NMDAR dysfunction through resultant developmental changes (6), suggesting that some of the features of systemic NMDAR hypofunction can be captured with cell-type specific effects. Recently inhibitory interneuronal dysfunction, particularly in the prefrontal circuit have emerged as a potential mechanism causing aspects of the schizophrenia phenotype (72, 73), which is further supported by computational models of prefrontal cortex functions (74).

Invasive recordings in the mouse prefrontal cortex suggest that in the microcircuit, the overall effect of NMDAR-transmission on inhibitory interneurons outweigh those directly on excitatory pyramidal cells: Specific NMDAR blockers cause a disinhibition effect by reducing the NMDAR-dependent activity of GABAergic interneurons and thus reducing inhibitory drive (53). In humans, proton magnetic resonance spectroscopy also suggests that there is a paradoxical increase in anterior cingulate glutamate in response to NMDAR blockade with ketamine (75).

Thus, the DCM findings presented here provide functional evidence for ketamine to alter human dynamic cortical networks in the way suggested by the animal model findings. Our study provides evidence for a regionally selective disinhibition in response to NMDAR blockade, and thus further supports NMDAR hypofunction specific to GABAergic interneurons as a mechanism underlying ketamine-induced cortical abnormalities.

## METHODS

### Subjects

For this study, N=18 healthy male volunteers were recruited from the local university through advertisement. All subjects gave fully informed, written consent prior to participation in the study and the study was approved by the University of Lübeck Research Ethics Committee. Subjects were compensated for the time they contributed to the research study. Table 1 presents subject details.

All subjects answered a basic clinical and a more focussed psychiatric (SCL-90-R®) questionnaire and underwent a routine clinical examination (including ECG, auscultation, and blood pressure measurements). Subjects with pre-existing neurological, psychiatric or cardiac conditions, those with a family history of psychotic illness or epilepsy, and subjects on any regular medication were excluded from participation. Further exclusion criteria were left-handedness, smoking, and the regular use of recreational drugs. Subjects were invited for two study sessions >3 weeks apart to participate both in the placebo, and the ketamine arm of the study.

The experiment followed a randomised, placebo-controlled, double-blind cross-over design: Subjects were allocated to either ketamine-or placebo-first groups, which was counter-balanced across subjects (9 in each group). Subjects and the researcher supervising EEG recording collection were blinded to the trial condition. However, because the higher ketamine dose was poorly tolerated by some subjects, the ketamine dose was reduced for subsequent data collection – the doses received by each subject are included in Table 1.

### Stimuli and EEG recording

Electroencephalography was conducted using 20 electrodes (10-20 international system), placed using an EASYCAP electrode cap (www.easycap.de). Signals were recorded through a Compumedics Neuroscan© amplifier system. Auditory stimuli were pure tones presented in pseudorandom sequences: In the roving paradigm, sounds of the same frequency were repeated for an unpredictable number of repetitions (range 6-36), before either the frequency or the duration of individual stimuli changes (76). Sounds were presented at a volume of 80dB at frequencies ranging from 700-1200 Hz, and with an inter-stimulus interval of 400ms. Subjects were given an incidental reading task and were instructed to ignore the sounds.

Data were processed in an average referential montage, recorded at a sampling frequency of 500Hz and bandpass filtered to a frequency range of 0.1 – 80Hz. For each test-session, data were divided into-100ms to 300ms peristimulus epochs. The roving paradigm contains deviant stimuli (e.g. the first sound presented at a new frequency), and standard sounds (e.g. multiple repetitions of the same sound). Because the characteristics of deviants and standards are not fixed, but with each new sequence of tones the standards are re-learned, evoked response potentials can be quantified for the tone-by-tone transition from deviant to standard; as an initially novel sound is repeated and becomes a standard. Thus, we calculated average ERPs for the 1st (i.e., deviant D1), 2nd, 6th, and 36th (i.e., standards S2, S6, and S36) repetition of each tone. Baseline correction was performed based on 100-0ms peristimulus time segments only for D1, 250-300ms peristimulus time segments for S2; and 100-0ms & 250-300ms peristimulus time segments for S6 and S36 (to avoid including the large P3a component present at the end of D1 – and prior to S2 – in the baseline correction, see Fig 2).

### Dynamic causal modelling

Dynamic causal modelling (DCM) is a standard Bayesian technique to estimate the parameters of neural mass models of cortical activity from EEG measurements. Here, we apply a hierarchical (parametric empirical) Bayesian modelling approach to identify parameter changes across families of DCMs, in order to estimate the effects of ketamine on cortical processing of regular auditory sequences and their violations. All DCM analyses were performed using the free academic software SPM12 (http://www.fil.ion.ucl.ac.uk/spm/, Litvak et al. 2011), and custom code available online. (doi.org/10.5281/zenodo.570595).

*Identifying prior parameter distributions from DCM on grand mean ERP curves*

In order to produce the best fits for model inversion at the level of single subjects, a DCM was first fitted to the grand mean average of the ERPs in the placebo condition. This grand mean DCM allowed changes in all model parameters. ERPs to the 1st (i.e. deviant D1), 2nd, 6th and 36th (i.e. standards S2, S6, S36) exposure to the same sound were modelled as connectivity changes in auditory network connectivity as described previously (49), and detailed below.

A standard electromagnetic forward model was generated, linking channel-level observations with cortical activity based on a template three-shell cortical mesh in MNI space. The resulting lead-fields were used to reconstruct source ERP waveforms from six cortical locations taken from previously published literature (22, 46): left and right primary auditory cortex (A1), left and right superior temporal gyrus (STG), and left and right inferior frontal gyrus (IFG). The MNI coordinates used were: ltA1 [-42,-22, 7], rtA1 [46,-14, 8], ltSTG [-61,-32, 8], rtSTG [59,-25, 8], ltIFG [-46, 20, 8], rtIFG [46, 20, 8].

In DCM, ERPs are modelled as neural population responses in a hierarchical network of reciprocally coupled sources with recurrent self-connections. Differences between ERPs were assumed to arise from changes in synaptic coupling; altering forward, backward and intrinsic (self) connection strengths between conditions. In analogy to the existing literature, we modelled the effect of deviancy as coupling changes (i.e., short-term plasticity) between presentations of the same sound using two temporal basis functions (monophasic decay, and a phasic effect, Fig 2A). Model inversion using these basis functions estimate the linear combination of the two effects that best explains the observed data, thus modelling a broad range of plausible time courses (49).

The variational Bayes model inversion implemented in DCM provides both a measure for the log-model evidence (in the form of model free energy), and posterior densities for the parameter values. The expected parameter estimates (without the posterior covariances) from the grand average were used as prior expectations for the inversion of individual subjects in the second step of the DCM analysis.

*Individual model inversion and Bayesian model reduction to identify repetition effects*

Whole-scalp ERPs for each subject were extracted separately for the placebo and the ketamine condition, resulting in 36 separate sessions (18 subjects, 2 conditions) for DCM analysis. For each DCM, the full network as described above, consisting of six hierarchically coupled cortical sources, was then equipped with prior expectations derived from the grand mean model inversion. Model inversion yielded separately parameterised DCMs for each subject and drug-condition, as well as an estimate of the associated log-model evidence.

To test whether network changes – estimated under our model – for both the difference between deviant and standard responses (50) and the effect of repetition (49) replicate existing findings in the literature, we then performed Bayesian model reduction and implicit comparison. In other words, we identified the best combination of parameter changes that could explain the observed ERP responses: Bayesian model reduction uses posterior parameter densities and free energy estimates from a fall or parent DCM to estimate the model-evidence for a number of reduced or children DCMs, in which some combinations of parameters are fixed or do not allow condition specific variations. This approach replaces the previous standard approach of inverting several reduced DCMs separately and provides a computationally efficient alternative estimation of the log-likelihood across a large space of models (44, 52, 54). The model space was based on the previous literature (22, 50), distinguishing between repetition-related changes of forward connections, backward connections and intrinsic modulatory gain parameters across a set of 2x2x2 = 8 factorial models (Fig 2A).

Bayesian model comparison identified the reduced model that best explained the data for each subject. As each subject had a high model evidence for the same (winning) model (see Results), effects across the group were summarised using Bayesian parameter averages to quantify parametric changes in forward connections, backward connections and intrinsic modulatory gain across repetitions.

*Parametric empirical Bayes and ketamine effects*

DCM inversions at the first (session) level furnish posterior estimates of the repetition effects on connectivity independently for placebo and ketamine. To estimate systematic changes in model parameters with ketamine dose (none for placebo, low or high for the two dosages) within a Bayesian modelling framework, we applied a parametric empirical Bayesian (PEB) approach (78). In brief, PEB allows the Bayesian estimation of a general linear model explaining effects across individually inverted DCMs. This enables estimation of parametric random effects and inference about treatment effects that are common to all the individual DCMs. The second level model can be equipped with a number of different regressors, and estimation of the second level model provides both estimates of these second level DCM parameters, as well as the log-model evidence for the hierarchical model; thereby allowing Bayesian model comparison at the second level.

Here we use PEB to (i) perform Bayesian model comparison across reduced models, where only a limited set of extrinsic or intrinsic DCM parameters are used to explain the changes induced by ketamine, and (ii) quantify the parameter changes caused by ketamine in the winning model.

Each PEB model contained all DCMs inverted under placebo, and ketamine for each individual (i.e. 36 first level DCMs). Regressors used for the PEB comprised the effect of ketamine (0 for placebo, 1 for low dose ketamine, 2 for high dose ketamine), the group mean and subject or block effects. Each PEB model differed only in which synaptic parameters (see Table 2) were used to explain ketamine effects. The parameters we considered included time constants (describing the shape of the kernel which maps the presynaptic inputs onto the postsynaptic membrane potentials), intrinsic connectivity parameters (quantifying the synaptic weights linking populations within cortical sources), modulatory gain parameters (quantifying the self-inhibition or gain of superficial pyramidal population, additionally modulated by inputs it receives from other regions), and extrinsic connectivity parameters (quantifying the weights linking different cortical sources). Crucially, we examined an effect of ketamine on (intrinsic and extrinsic) connectivity *and repetition dependent effects* parameterised in terms of basis functions at the first level. In other words, we allowed for both a non-specific (main) effect of ketamine on coupling – and an (interaction) effect on plasticity (in DCM, these are usually referred to as A and B parameters – see Tab 1). Bayesian model reduction of these second level models provided estimates of their associated log-evidence, which was used to identify the winning second-level model.

The winning (reduced) model also provided estimates of the direction and size of the parameter changes induced by ketamine. Because these are estimated as posterior densities, they provide both an estimate of ketamine effects, and uncertainty around these effects, which can be used to estimate a Bayesian 95% confidence interval. Finally, in order to characterise the effects of the parameter changes induced by ketamine, we used the parameter estimates in simulation mode (i.e., in a forward model) to visualise their effects on the source-space ERPs. This forward model was based on the DCM fitted to the grand mean across the whole group.

## Acknowledgements

RER is funded by a Wellcome Trust Clinical Research Fellowship (106556/Z/14/Z). KJF is funded by a Wellcome Trust Principal Research Fellowship (088130/Z/09/Z).

## REFERENCES

1. Friston K, Brown HR, Siemerkus J, Stephan KE (2016) The dysconnection hypothesis (2016). Schizophr Res 176(2-3):83–94.

2. Scoriels L, et al. (2015) Behavioural and molecular endophenotypes in psychotic disorders reveal heritable abnormalities in glutamatergic neurotransmission. Transl Psychiatry 5(3):e540.

3. Wang X-J, Krystal JH (2014) Computational Psychiatry. Neuron 84(3):638–654.

4. Corlett PR, Honey GD, Krystal JH, Fletcher PC (2011) Glutamatergic Model Psychoses: Prediction Error, Learning, and Inference. Neuropsychopharmacology 36(1):294–315.

5. Krystal JH (1994) Subanesthetic Effects of the Noncompetitive NMDA Antagonist, Ketamine, in Humans. Arch Gen Psychiatry 51(3):199.

6. Nakazawa K, Jeevakumar V, Nakao K (2017) Spatial and temporal boundaries of NMDA receptor hypofunction leading to schizophrenia. npj Schizophr 3(1):7.

7. Guo X, et al. (2009) Reduced Expression of the NMDA Receptor-Interacting Protein SynGAP Causes Behavioral Abnormalities that Model Symptoms of Schizophrenia. Neuropsychopharmacology 34(7):1659–1672.

8. Javitt DC, Steinschneider M, Schroeder CE, Arezzo JC (1996) Role of cortical N-methyl-D-aspartate receptors in auditory sensory memory and mismatch negativity generation: implications for schizophrenia. Proc Natl Acad Sci 93(21):11962–11967.

9. Urs NM, Peterson SM, Caron MG (2017) New Concepts in Dopamine D2 Receptor Biased Signaling and Implications for Schizophrenia Therapy. Biol Psychiatry 81(1):78–85.

10. Catts VS, Lai YL, Weickert CS, Weickert TW, Catts S V. (2016) A quantitative review of the postmortem evidence for decreased cortical N-methyl-d-aspartate receptor expression levels in schizophrenia: How can we link molecular abnormalities to mismatch negativity deficits? Biol Psychol 116:57–67.

11. Glausier JR, Lewis DA (2013) Dendritic spine pathology in schizophrenia. Neuroscience 251:90–107.

12. Takahashi N, Sakurai T, Davis KL, Buxbaum JD (2011) Linking oligodendrocyte and myelin dysfunction to neurocircuitry abnormalities in schizophrenia. Prog Neurobiol 93(1):13–24.

13. Hu M, et al. (2013) Decreased left middle temporal gyrus volume in antipsychotic drug-naive, first-episode schizophrenia patients and their healthy unaffected siblings. Schizophr Res 144(1-3):37–42.

14. Stephan KE, Baldeweg T, Friston KJ (2006) Synaptic Plasticity and Dysconnection in Schizophrenia. Biol Psychiatry 59(10):929–939.

15. Anticevic A, et al. (2012) NMDA receptor function in large-scale anticorrelated neural systems with implications for cognition and schizophrenia. Proc Natl Acad Sci 109(41):16720–16725.

16. Moran RJ, Symmonds M, Stephan KE, Friston KJ, Dolan RJ (2011) An In Vivo Assay of Synaptic Function Mediating Human Cognition. Curr Biol 21(15):1320–1325.

17. Ranlund S, et al. (2016) Impaired prefrontal synaptic gain in people with psychosis and their relatives during the mismatch negativity. Hum Brain Mapp 37(1):351–365.

18. Fogelson N, Litvak V, Peled A, Fernandez-del-Olmo M, Friston K (2014) The functional anatomy of schizophrenia: A dynamic causal modeling study of predictive coding. Schizophr Res 158(1-3):204–212.

19. Kömek K, Bard Ermentrout G, Walker CP, Cho RY (2012) Dopamine and gamma band synchrony in schizophrenia-insights from computational and empirical studies. Eur J Neurosci 36(2):2146–2155.

20. Umbricht D, Krljes S (2005) Mismatch negativity in schizophrenia: a meta-analysis. Schizophr Res 76(1):1–23.

21. Wynn JK, Sugar C, Horan WP, Kern R, Green MF (2010) Mismatch Negativity, Social Cognition, and Functioning in Schizophrenia Patients. Biol Psychiatry 67(10):940–947.

22. Phillips HN, Blenkmann A, Hughes LE, Bekinschtein T a., Rowe JB (2015) Hierarchical Organization of Frontotemporal Networks for the Prediction of Stimuli across Multiple Dimensions. J Neurosci 35(25):9255–9264.

23. Cheour M, H.T. Leppänen P, Kraus N (2000) Mismatch negativity (MMN) as a tool for investigating auditory discrimination and sensory memory in infants and children. Clin Neurophysiol 111(1):4–16.

24. Todd J, et al. (2014) Mismatch negativity (MMN) to pitch change is susceptible to order-dependent bias. Front Neurosci 8(8 JUN):1–9.

25. El Karoui I, et al. (2015) Event-Related Potential, Time-frequency, and Functional Connectivity Facets of Local and Global Auditory Novelty Processing: An Intracranial Study in Humans. Cereb Cortex 25(11):4203–4212.

26. Baldeweg T, Hirsch SR (2015) Mismatch negativity indexes illness-specific impairments of cortical plasticity in schizophrenia: A comparison with bipolar disorder and Alzheimer’s disease. Int J Psychophysiol 95(2):145–155.

27. Auksztulewicz R, Friston K (2015) Attentional Enhancement of Auditory Mismatch Responses: a DCM/MEG Study. Cereb Cortex 25(11):4273–4283.

28. Auksztulewicz R, Friston K (2016) Repetition suppression and its contextual determinants in predictive coding. Cortex 80:125–140.

29. Ostwald D, et al. (2012) Evidence for neural encoding of Bayesian surprise in human somatosensation. Neuroimage 62(1):177–188.

30. Astikainen P, Lillstrang E, Ruusuvirta T (2008) Visual mismatch negativity for changes in orientation-a sensory memory-dependent response. Eur J Neurosci 28(11):2319–2324.

31. PulverMüller F, Assadollahi R (2007) Grammar or Serial Order?: Discrete Combinatorial Brain Mechanisms Reflected by the Syntactic Mismatch Negativity. J Cogn Neurosci 19(6):971–980.

32. Friston K, Kiebel S (2009) Predictive coding under the free-energy principle. Philos Trans R Soc B Biol Sci 364(1521):1211–1221.

33. Schröger E, et al. (2014) Predictive Regularity Representations in Violation Detection and Auditory Stream Segregation: From Conceptual to Computational Models. Brain Topogr 27(4):565–577.

34. Dayan P, Hinton GE, Neal RM, Zemel RS (1995) The Helmholtz Machine. Neural Comput 7(5):889–904.

35. Friston K (2005) A theory of cortical responses. Philos Trans R Soc B Biol Sci 360(1456):815–836.

36. Jaaskelainen IP, et al. (2004) Human posterior auditory cortex gates novel sounds to consciousness. Proc Natl Acad Sci 101(17):6809–6814.

37. Näätänen R, Winkler I (1999) The concept of auditory stimulus representation in cognitive neuroscience. Psychol Bull 125(6):826–859.

38. Todd J, Petherbridge A, Speirs B, Provost A, Paton B (2017) Time as context: The influence of hierarchical patterning on sensory inference. Schizophr Res. doi:10.1016/j.schres.2017.03.033.

39. Ylinen S, et al. (2016) Predictive coding of phonological rules in auditory cortex: A mismatch negativity study. Brain Lang 162:72–80.

40. Wacongne C, Changeux J-P, Dehaene S (2012) A Neuronal Model of Predictive Coding Accounting for the Mismatch Negativity. J Neurosci 32(11):3665–3678.

41. Bastos AM, et al. (2012) Canonical Microcircuits for Predictive Coding. Neuron 76(4):695–711.

42. Moran R, Pinotsis DA, Friston K (2013) Neural masses and fields in dynamic causal modeling. Front Comput Neurosci 7(May):57.

43. Friston KJ, Harrison L, Penny W (2003) Dynamic causal modelling. Neuroimage 19(4):1273–1302.

44. Kiebel SJ, Garrido MI, Moran R, Chen C-C, Friston KJ (2009) Dynamic causal modeling for EEG and MEG. Hum Brain Mapp 30(6):1866–1876.

45. Cooray G, Garrido MI, Hyllienmark L, Brismar T (2014) A mechanistic model of mismatch negativity in the ageing brain. Clin Neurophysiol 125(9):1774–1782.

46. Garrido MI, et al. (2008) The functional anatomy of the MMN: A DCM study of the roving paradigm. Neuroimage 42(2):936–944.

47. Cooray GK, Garrido MI, Brismar T, Hyllienmark L (2016) The maturation of mismatch negativity networks in normal adolescence. Clin Neurophysiol 127(1):520–529.

48. Dima D, Frangou S, Burge L, Braeutigam S, James AC (2012) Abnormal intrinsic and extrinsic connectivity within the magnetic mismatch negativity brain network in schizophrenia: A preliminary study. Schizophr Res 135(1-3):23–27.

49. Garrido MI, et al. (2009) Repetition suppression and plasticity in the human brain. Neuroimage 48(1):269–279.

50. Garrido MI, Kilner JM, Kiebel SJ, Friston KJ (2008) Dynamic Causal Modeling of the Response to Frequency Deviants. J Neurophysiol 101(5):2620–2631.

51. Schmidt A, et al. (2013) Modeling Ketamine Effects on Synaptic Plasticity During the Mismatch Negativity. Cereb Cortex 23(10):2394–2406.

52. Friston K, Zeidman P, Litvak V (2015) Empirical Bayes for DCM: A Group Inversion Scheme. Front Syst Neurosci 9(November):1–10.

53. Homayoun H, Moghaddam B (2007) NMDA Receptor Hypofunction Produces Opposite Effects on Prefrontal Cortex Interneurons and Pyramidal Neurons. J Neurosci 27(43):11496–11500.

54. Litvak V, Garrido M, Zeidman P, Friston K (2015) Empirical Bayes for Group (DCM) Studies: A Reproducibility Study. Front Hum Neurosci 9(Dcm):1–12.

55. Rosier A, Arckens L, Orban G, Vandesande F (1993) Laminar distribution of NMDA receptors in cat and monkey visual cortex visualized by [3H]-MK-801 binding. J Comp Neurol 335(3):369–380.

56. Moreau AW, Kullmann DM (2013) NMDA receptor-dependent function and plasticity in inhibitory circuits. Neuropharmacology 74:23–31.

57. Bond T, Durrant S, O’Hare L, Turner D, Sen-Bhattacharya B (2014) Studying the effects of thalamic interneurons in a thalamocortical neural mass model. BMC Neurosci 15(Suppl 1):P219.

58. Bhattacharya B Sen, Bond TP, O’Hare L, Turner D, Durrant SJ (2016) Causal Role of Thalamic Interneurons in Brain State Transitions: A Study Using a Neural Mass Model Implementing Synaptic Kinetics. Front Comput Neurosci 10(November):1–18.

59. Jirsa VK, Stacey WC, Quilichini PP, Ivanov AI, Bernard C (2014) On the nature of seizure dynamics. Brain 137(8):2210–2230.

60. Wendling F, Benquet P, Bartolomei F, Jirsa V (2016) Computational models of epileptiform activity. J Neurosci Methods 260:233–251.

61. Jentsch JD, Roth RH (1999) The Neuropsychopharmacology of Phencyclidine?: From NMDA Receptor Hypofunction to the Dopamine Hypothesis of Schizophrenia. Neuropsychopharmacology 20(3):201–225.

62. Geyer MA, Krebs-Thomson K, Braff DL, Swerdlow NR (2001) Pharmacological studies of prepulse inhibition models of sensorimotor gating deficits in schizophrenia: A decade in review. Psychopharmacology (Berl) 156(2-3):117–154.

63. Murray JD, et al. (2014) A hierarchy of intrinsic timescales across primate cortex. Nat Neurosci 17(12):1661–1663.

64. Chaudhuri R, Bernacchia A, Wang X-J (2014) A diversity of localized timescales in network activity. Elife 3:e01239.

65. Kiebel SJ, Daunizeau J, Friston KJ (2008) A Hierarchy of Time-Scales and the Brain. PLoS Comput Biol 4(11):e1000209.

66. Ulanovsky N (2004) Multiple Time Scales of Adaptation in Auditory Cortex Neurons. J Neurosci 24(46):10440–10453.

67. Nelken I, Fishbach A, Las L, Ulanovsky N, Farkas D (2003) Primary auditory cortex of cats: feature detection or something else? Biol Cybern 89(5):397–406.

68. Bastos AM, et al. (2015) A DCM study of spectral asymmetries in feedforward and feedback connections between visual areas V1 and V4 in the monkey. Neuroimage 108:460–475.

69. Muthukumaraswamy SD, et al. (2015) Evidence that Subanesthetic Doses of Ketamine Cause Sustained Disruptions of NMDA and AMPA-Mediated Frontoparietal Connectivity in Humans. J Neurosci 35(33):11694–11706.

70. Moran RJ, et al. (2015) Losing Control Under Ketamine: Suppressed Cortico-Hippocampal Drive Following Acute Ketamine in Rats. Neuropsychopharmacology 40(2):268–277.

71. Bygrave A, et al. (2016) Knockout of NMDA-receptors from parvalbumin interneurons sensitizes to schizophrenia-related deficits induced by MK-801. Transl Psychiatry 6(4):e778.

72. Cohen SM, Tsien RW, Goff DC, Halassa MM (2015) The impact of NMDA receptor hypofunction on GABAergic neurons in the pathophysiology of schizophrenia. Schizophr Res 167(1-3):98–107.

73. Lewis D a, Hashimoto T, Volk DW (2005) Cortical inhibitory neurons and schizophrenia. Nat Rev Neurosci 6(4):312–324.

74. Murray JD, et al. (2014) Linking Microcircuit Dysfunction to Cognitive Impairment: Effects of Disinhibition Associated with Schizophrenia in a Cortical Working Memory Model. Cereb Cortex 24(4):859–872.

75. Stone JM, et al. (2012) Ketamine effects on brain GABA and glutamate levels with 1H-MRS: relationship to ketamine-induced psychopathology. Mol Psychiatry 17(7):664–665.

76. Haenschel C (2005) Event-Related Brain Potential Correlates of Human Auditory Sensory Memory-Trace Formation. J Neurosci 25(45):10494–10501.

77. Litvak V, et al. (2011) EEG and MEG Data Analysis in SPM8. Comput Intell Neurosci 2011:1–32.

78. Friston KJ, et al. (2016) Bayesian model reduction and empirical Bayes for group (DCM) studies. Neuroimage 128:413–431.

